# Extended depth of focus multiphoton microscopy via incoherent pulse splitting

**DOI:** 10.1101/2020.03.27.012260

**Authors:** Bingying Chen, Tonmoy Chakraborty, Stephan Daetwyler, James D. Manton, Kevin Dean, Reto Fiolka

## Abstract

We present a phase mask that can be easily added to any multi-photon raster scanning microscope to extend the depth of focus five-fold at a small loss in lateral resolution. The method is designed for ultrafast laser pulses or other light-sources featuring a low coherence length. In contrast to other methods of focus extension, our approach uniquely combines low complexity, high light-throughput and multicolor capability. We characterize the point-spread function in a two-photon microscope and demonstrate fluorescence imaging of GFP labeled neurons in fixed brain samples as imaged with conventional and extended depth of focus two-photon microscopy.

## 1. Introduction

Imaging of large volumes with raster scanning microscopes can be very time consuming as they acquire each voxel serially. This is particularly true if high NA objectives are employed that provide a high lateral resolution and a small depth of focus. To capture volumetric imaging data exceeding such a small depth, many planes have to be imaged by z-stepping each focal plane, which limits the achievable imaging speed considerably. Instead, if axial image information can be sacrificed, it can be beneficial to extend the depth of focus such that the volume to be imaged can be acquired in one lateral scan. In other words, the volumetric information is projected into a single 2D image. As such, extended depth of focus (EDF) imaging can be attractive to image sparsely populated structures rapidly and has found promising applications in functional imaging of neuronal activity [1,2].

Various beam shaping methods have been successfully employed for EDF imaging, which involve different amplitude, phase or combined filters in the pupil plane [3–5], and in some cases also feature specialized optical elements such as axicons [6,7]. Their implementation typically requires significant changes to a conventional microscope [1]. Further, if phase modulation is used, the associated phase masks or holograms are typically wavelength dependent, which prevents, or at least complicates, multi-color imaging.

A conceptually different approach was presented by Abrahamsson and Gustafsson [8], which uses an incoherent superposition as the means of creating an EDF beam: Here, the pupil of an objective is segmented into multiple annuli, which are made incoherent to each other. The exact relative phase between the different annuli does not matter if the resulting beams are mutually incoherent. This concept was designed for increasing the depth in the detection path of a fluorescent widefield microscope, exploiting the short coherence length of fluorescence. However, we reasoned that this concept should also be applicable to beam shaping of ultrafast laser sources, which can feature a similarly short coherence length.

In this manuscript, we present numerical simulations and experimental measurements of two-photon EDF microscopy using this incoherent annular phase mask. In addition, we highlight its potential for imaging applications in neuroscience by imaging GFP-labeled neurons in a fixed brain slice.

## 2. Method

### 2.1 Theory

The phase mask, which is acting in a Fourier plane of the imaging system, consists of multiple concentric glass disks (Figure 1a). An ultrafast laser pulse (typical pulse length in the tens of microns) is split by the phase mask into different annular beamlets (Figure 1b), each of which is time-delayed and form a focus in the front focal plane of the objective at slightly different arrival times. In the time average, the resulting EDF focus is the incoherent superposition of all individual foci (Figure 1c), which led us term the technique incoherent pulse splitting. As shown in previous work using such a phase mask[8], for processes that scale linearly with intensity, some strong sidelobe structures will surround the main lobe of this EDF focus. In our case, we consider two-photon absorption, which dampens sidelobe structures and yields a smooth, gradual decay into the background (Figure 1c).

**Fig. 1.**
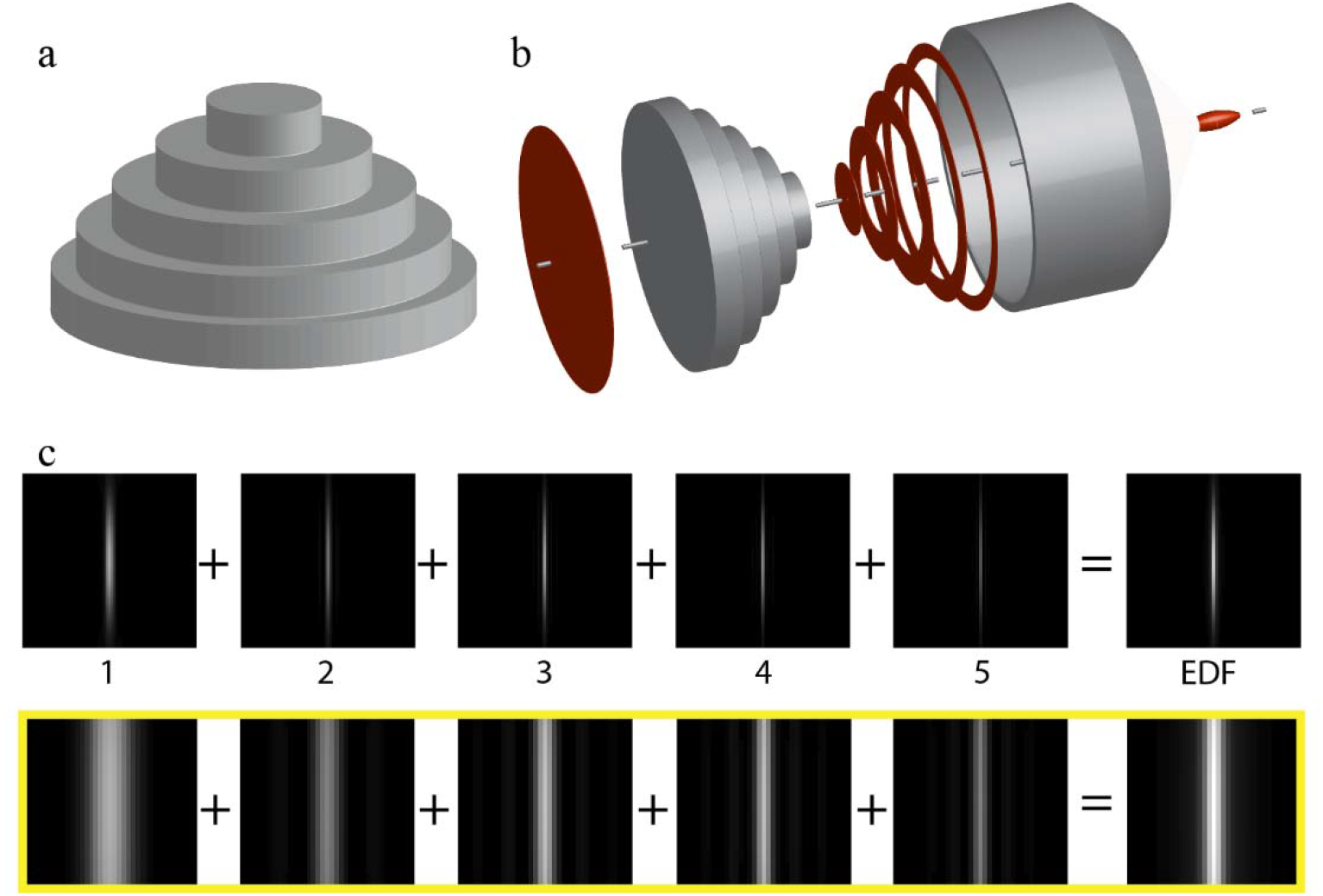
Principle of extended depth of focus for ultrafast laser pulses. (a) Schematic of the phase mask consisting of multiple glass disks of varying diameter. (b) Illustration of the working principle. An ultrafast laser pulse (red disk) is incident on the phase mask. Portions of the pulse travelling through the different annular zones get time delayed into separate beamlets (red annuli). Each beamlet forms a focus at the front focal plane of the objective at a different time. (c) Numerical simulations of the squared intensity of the laser foci produced by the different beamlets. The effective PSF is the incoherent sum of all five foci. Yellow box shows the zoomed in view (centered on the bigger views in the row above) of the PSFs.

Previous work suggested that the extension of the depth of focus scales with the number of annular zones[8]. Thus, for a phase mask with five zones, a roughly fivefold extension is expected. Using a numerical aperture >1 generates point spread functions with axial widths in the range of a single micrometer. Even with a fivefold extension by our phase mask, this would still be a rather modest depth of focus. We therefore reason that using a medium numerical aperture for excitation (in the range of 0.5-0.7) would still provide sufficient lateral resolution for many imaging applications in neuroscience while offering a more sizeable depth of focus when combined with our phase mask (e.g., 15-20 microns).

### 2.2 Simulations

Numerical simulations to compute the electromagnetic field in the front focal plane of an objective were performed using the Debye theory [9]. In short, the simulation calculated the 3D electromagnetic field in the focal volume for each annular pupil zone individually. Each electric field was converted to an intensity by squaring the modulus of the field and then further converted to a multiphoton PSF via another squaring operation. The resulting PSFs were then summed to form the overall EDF PSF. A conventional PSF, using the same maximal numerical aperture, is also calculated to enable comparison. The simulation results predict an extension of the depth of focus that scales with the number of annular zones, consistent with earlier reports using this mask for depth of focus extension in linear fluorescence imaging.

In general, the numerical apertures (NA) of the annuli can be calculated using the following procedure. First, calculate the central zone NA which produces a Gaussian-like PSF of the desired depth-of-field, where the depth-of-field *d* for a given wavelength λ and a given refractive index n is determined by the following equation:

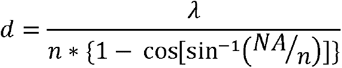

Next, set the first annular zone to have an inner NA, NA_in_, equivalent to the NA just calculated for the Gaussian-like PSF, and an outer NA, NA_out_, such that the Bessel-like PSF has a depth of field, given by

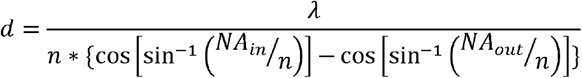

that matches the desired depth-of-field. Repeat this outer NA calculation process, using the previous outer NA as the inner NA of the next zone until all annuli are considered or the maximum available NA of the lens is used.

Here, as we wanted to match the physical parameters of the mask used for our experimental results, we instead converted the annular diameter of the mask (1.8 mm, 2.5 mm, 3.1 mm, 3.6 mm, 4.0 mm) to NAs, taking into account the magnification between mask and pupil, and calculated the corresponding electric fields. We chose, by selecting the magnification of the pupil relay, to use a maximum NA of 0.67, which yielded in the simulation a lateral resolution in the range of 500nm and an axial Full-Width Half-Maximum (FWHM) of ~16 microns (Figure 2 a). Compared to a conventional laser focus using this NA, an axial extension of the depth of focus of ~5 is achieved (Figure 2b). The simulation results suggest a 13% increase of the lateral Full Width Half Maximum (FWHM) of the EDF beam compared to a conventional laser focus (Figure 2c).

**Fig. 2.**
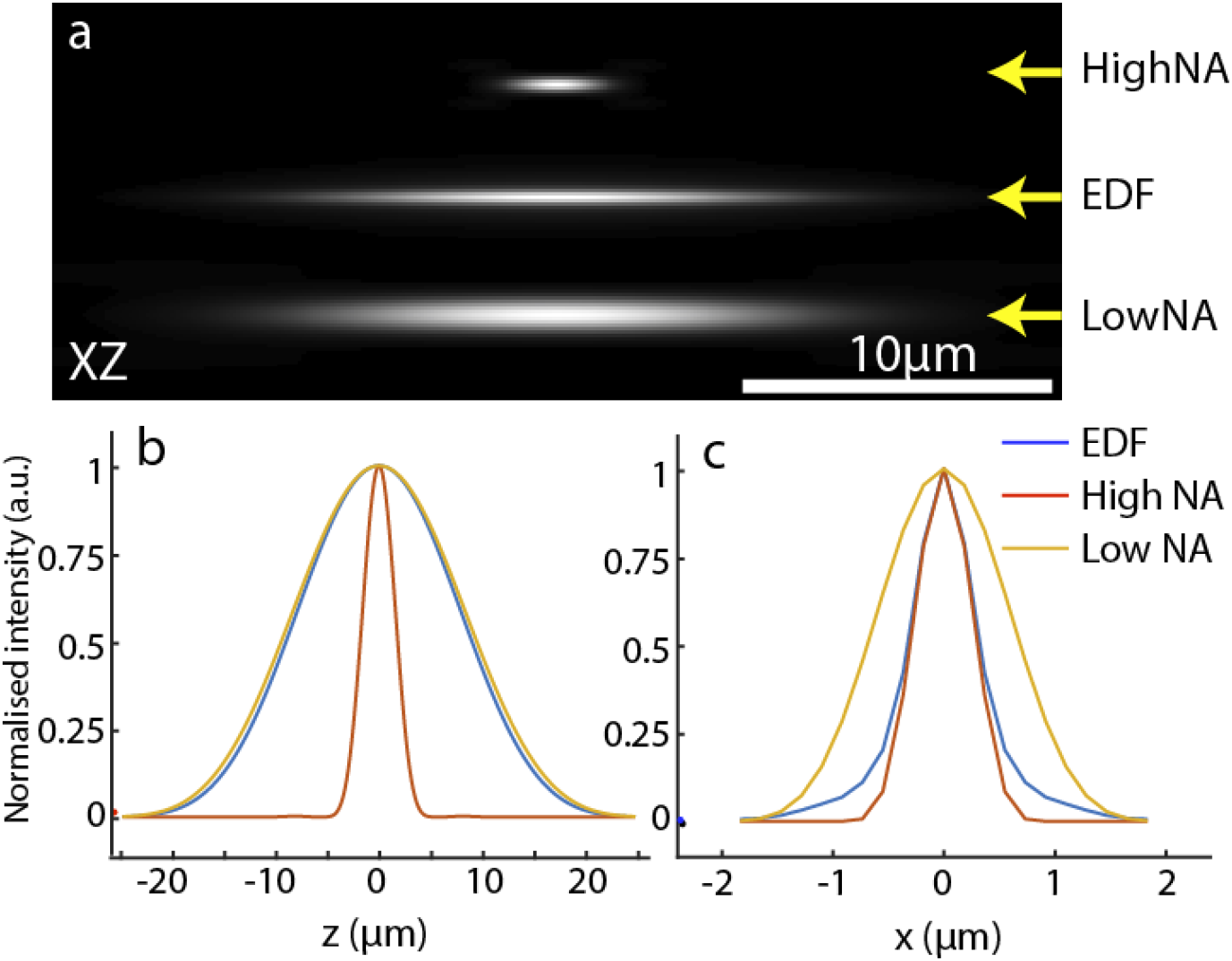
Numerical simulation of point spread functions (PSF). (a), from top to bottom: conventional PSF using an upper NA of 0.67 and 900nm wavelength. Middle: EDF PSF. Bottom: low NA PSF to match axial width of EDF PSF. (b-c) Axial and lateral cross-sections through the PSFs shown above.

### 2.3 Experimental setup

Our setup is based on a homebuilt two-photon raster scanning microscope as shown in Figure 3. Ultrafast laser pulses (900nm wavelength, ~150fs pulse lengths) from a Ti:Sapphire oscillator (Chameleon Vision, Coherent Inc) pass through a Pockel cell (Model 350-80-2, Conoptics) for power modulation and are expanded 6-fold by a Galilean telescope (AC254-50-B, AC254-300-B Thorlabs) and spatially cleaned up with a pinhole (PH-100, Newport). The expanded laser beam is incident on the phase mask (part Nr. A12802-35-040 Hamamatsu Photonics), and an iris was used to adjust the beam diameter to the size of the mask. The same iris was also used to create a low NA excitation beam. The phase mask is conjugated by a telescope onto the first galvo mirror (6215H, Cambridge technology). The galvo mirror is conjugated with a telescope consisting of two Plössl lenses (F-75 mm, built from two AC508-150-C each, Thorlabs) onto the second galvo mirror (6215H, Cambridge technology). A scan lens (SL50-2P2, Thorlabs) and tube lens (TL200-2P2, Thorlabs) are used to conjugate the galvos and the phase mask onto the pupil plane of the objective (XLPLN25XWMP2, NA 1.05 / 25X Olympus). The beam magnification was chosen such that an NA of 0.67 was used for excitation, i.e. the beam underfilled the pupil of the objective. A further experiment using a higher NA of the illumination objective (0.9) was performed using a different telescope to relay the phase mask onto the galvo mirrors (supplementary Note 1 and supplementary Figure 1).

**Fig. 3.**
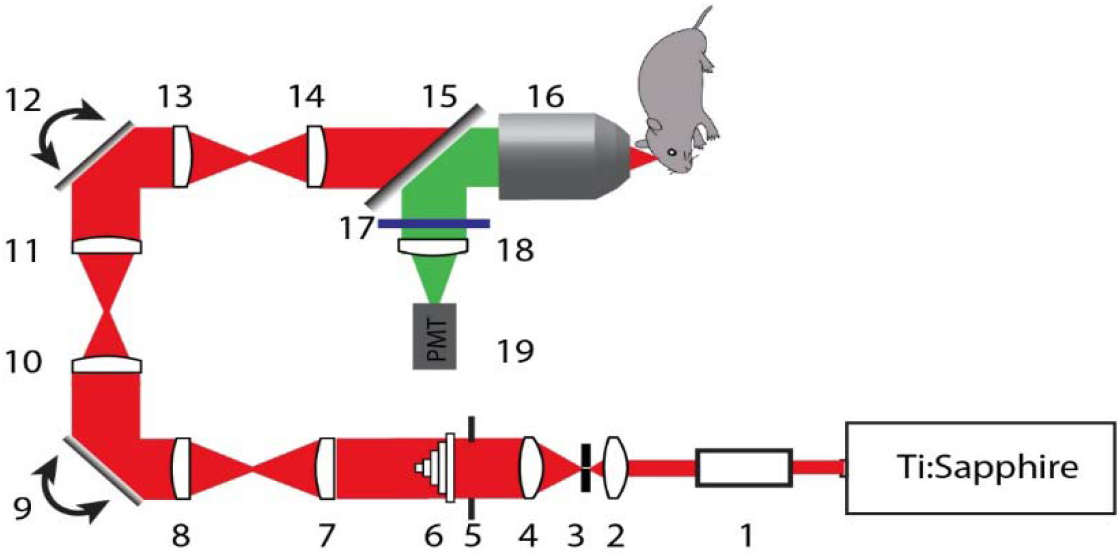
Schematic diagram of the experimental setup. 1, Pockel cell, 350-80-2, Conoptics; 2, AC254-50-B, Thorlabs; 3, Pinhole; 4, AC254-300-B; 5,Iris; 6, Phase mask A12802-35-040; 7 AC254-125-B; 8, AC254-75-B; 9,12, Galvo; 10, 11, Plössl lens; 13, Scan Lens, SL50-2P2; 14, Tube Lens, ,TL200-2P2; 15, Dichroic Mirror, FF735-Di02-50.8-D, Semrock; 16, Objective, XLPLN25XWMP2, Olympus; 17, Filters, FF02-694/sp-25, FF01-527/70-25, Semrock;18, AC254-45-A; 19, PMT, H7422-40, Hammamatsu

Fluorescence light was collected through the same objective, reflected by a dichromatic mirror (FF735-Di02-50.8-D, Semrock), filtered by a short-pass filter (FF02-694/sp-25, Semrock) and a bandpass filter (FF01-527/70-25, Semrock) and focused onto a PMT (H7422-40, Hammamatsu Inc). The signals from the PMT were amplified and filtered with an amplifier (DLPCA-200, Femto) and then read in with a DAQ card (PXIe-6341, X Series, National Instruments). The microscope was controlled using Scanimage (Vidrio Technologies, LLC, VA).

## 3. Results

### 3.1 Point spread function measurements

To measure the point spread functions (PSF) of our two-photon EDF system, we acquired 3D stacks of 200 nm green fluorescent beads (Polysciences, PA). Figure 4a shows the diffraction limited PSF obtained without the EDF phase mask using an NA of 0.67 for excitation (top), the PSF obtained using the phase mask (middle), and a PSF when the iris was closed such that only the innermost zone of the EDF mask contributed to the PSF (bottom). The latter represents a low NA Gaussian laser focus, which has the same confocal parameter as the EDF PSF (as can be seen in the axial line profiles). The axial FWHM is 16.83 ± 0.29 μm (mean and standard deviation, n=5) for the EDF PSF and 2.90 ± 0.08 μm for the PSF without the phase mask, which represents an axial extension by roughly a factor 5.8. In the lateral beam profile, one can see that the EDF PSF gets about 32% wider, increasing the width of the diffraction limited PSF of 0.53 ± 0.03 μm to 0.71 ± 0.02 μm. If using a low NA beam to match the axial FWHM of the EDF PSF, the width increases to 1.10 ± 0.06 μm.

**Fig. 4.**
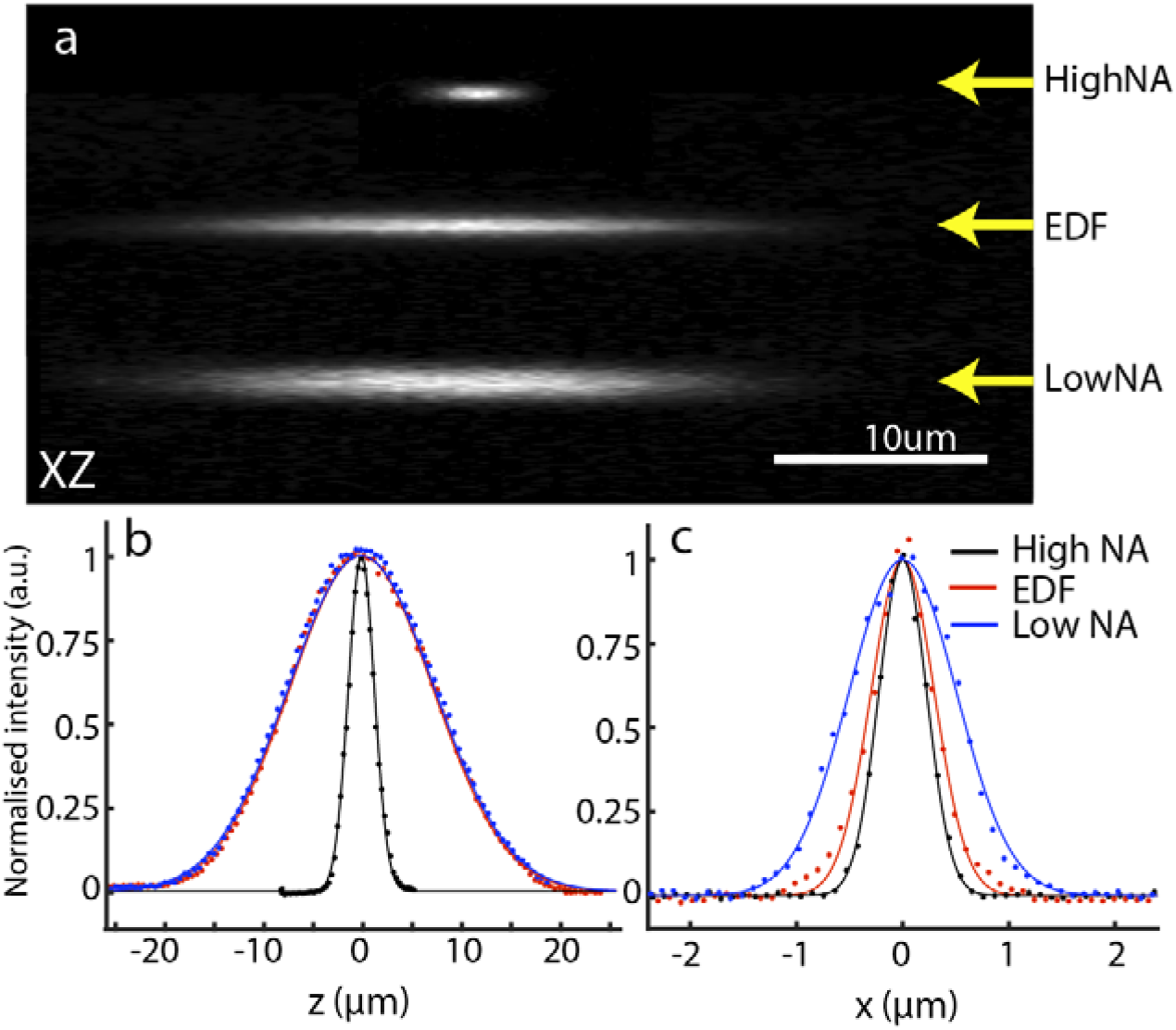
Point spread functions measured with 200 nm green fluorescent beads. (a), from top to bottom: PSF for laser focus using full NA of 0.67 and no EDF mask. PSF using EDF phase mask. Low NA PSF corresponding to the innermost zone of the EDF mask. (b-c) Axial and lateral profiles through the point spread functions. Black curve corresponds to full NA PSF without EDF mask, red curve corresponds to PSF using the EDF mask and blue curve corresponds to low NA laser focus.

### 3.2 Neuron imaging in fixed tissues

To demonstrate the potential of this EDF mask for neuroscience imaging applications, we imaged GFP labeled neurons in a fixed mouse brain tissue. For fixation, a Thy1-GFP mouse was transcardially perfused first with cold PBS, then with 4% PFA in PBS as described previously [10]. The brain was isolated and further fixed in 4% PFA in PBS overnight at 4°C. We first imaged a volume with the EDF mask as a single 2D image (Figure 5a). Next, we removed the EDF mask and acquired a conventional 3D image stack encompassing a 22.5-micron deep volume using conventional z-stepping by moving the sample. Figure 5b shows this volume color coded for depth. Close examination of a selected region (Figure 5 c-d) revealed close correspondence between the lateral features that can be seen in the EDF image and in the conventional 3D stack. Figure 5 e-g further shows individual z-planes from the conventional z-stack. Multiple cell bodies could only be seen in a specific z-plane, but not in others. All these cell bodies can be clearly seen in the single image frame acquired with the EDF phase mask. We note that our conventional z-stack fitted best to the EDF data when using a range of 22.5 microns, which is larger than the axial FWHM of the EDF PSF (~16 microns). We explain this discrepancy by the fact that in the EDF mode, we apparently still generated some useful signal outside of the full width half maximum range of the PSF.

**Fig. 5.**
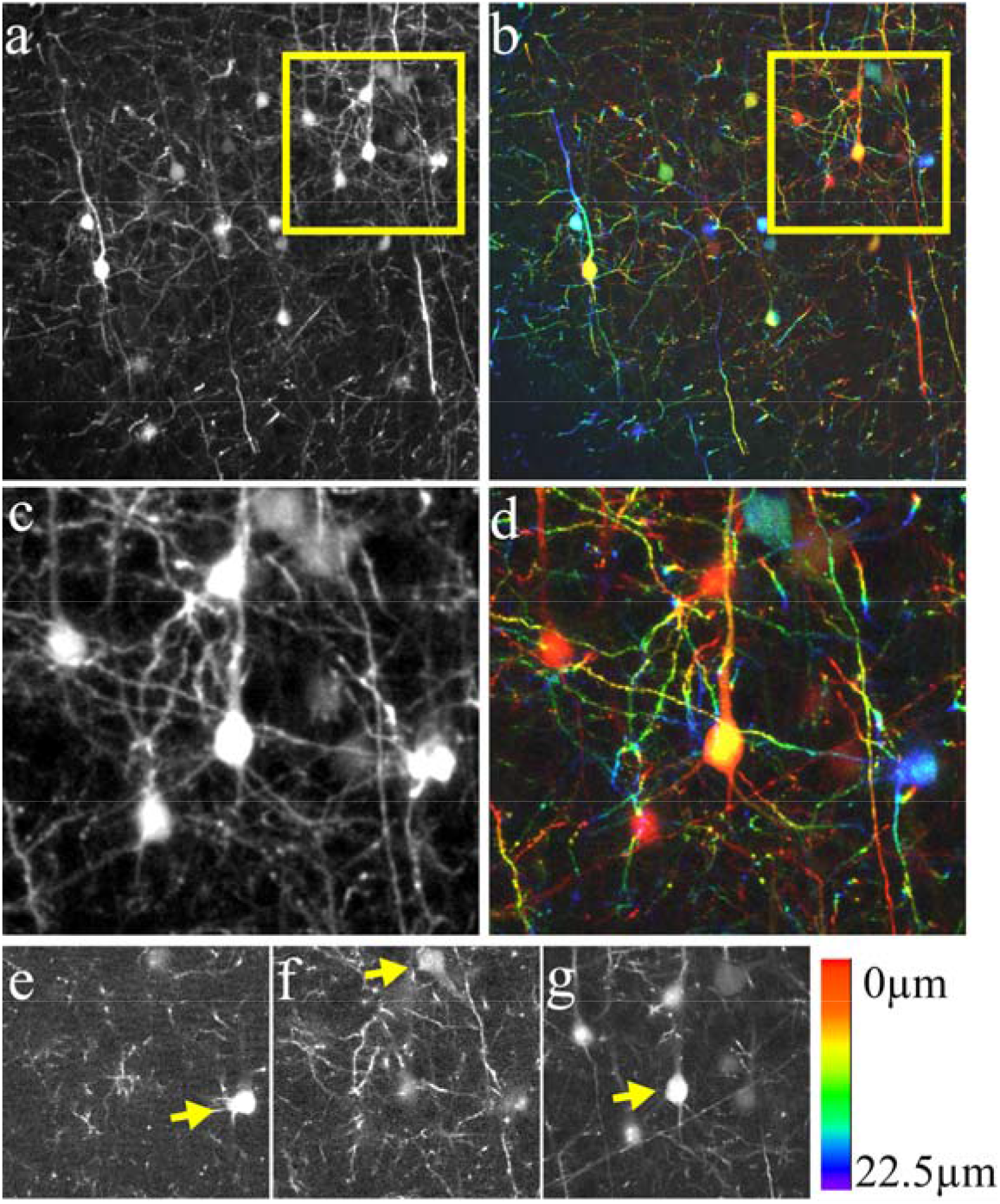
Volumetric two-photon imaging of Thy1-GFP labelled neurons in fixed brain tissue using either the EDF phase mask or conventional two-photon laser scanning with z-stepping. (a) single frame acquired with the EDF phase mask. (b) projection of a 3D volume acquired with conventional laser scanning over 16 z-planes spanning a range of 22.5 microns. Color encoded z-position. (c) magnified view of the boxed region of panel (a). (d) magnified region of the same area of panel b. (e-f) Individual z-planes from the region shown in panel d at 22.5 um, 10.5 um and 0 um depth. Arrows mark selected cell bodies that appear at these distinct depths.

## 4. Discussion

We have presented a simple way to extend the depth of focus in multiphoton microscopy, using a layered phase mask previously developed for fluorescence widefield imaging. We showed that this mask design can be advantageously used for extending the nonlinear focus in two-photon microscopy. The numerical and experimental results demonstrated that a five-fold extension in focal depth was feasible, without incurring sidelobes and with only a moderate lateral resolution loss. We note that while the simulation predicted a lateral resolution loss in the order of 13% in practice we measured a larger broadening of 32%. We also observed that with a larger maximum NA, the discrepancy between simulation and experiment increased (Supplementary Figure 1). We interpret this as a sign that our optical train may still have residual aberrations that do not manifest themselves as severely at a lower numerical aperture.

By imaging fixed brain samples we have demonstrated that the method delivered faithful images that offered comparable image details as the projection of conventionally acquired 3D image data. As such, we see wide applications for rapid volumetric imaging where multiple objects at different depths must be monitored simultaneously (e.g. calcium or voltage sensing in neurons). Also, the increased depth of focus could be helpful in scenarios of significant sample motion, as a much larger z-range remains within focus.

The method works on the principle that the time delay between the individual zones exceeds the pulse duration of the ultrafast laser pulse. As such, the method is broadly applicable to a wide range of pulsed laser sources (e.g. Q-switching or mode-locking) and insensitive to changes in the pulse wavelength. This is in stark contrast to methods that rely on phase masks that are precisely tailored to precise phase shifts for a specific wavelength.

Compared to two-photon raster scanning with Bessel beams, our method promises much higher light-throughput and reduced complexity. In fact, we showed that the mask can be directly placed in front of the galvanometric mirror (supplementary Note 2 and supplementary Figure 2). As such, any existing two-photon raster scanning microscope can be easily retrofitted without adding any other optical components than the mask. In contrast, published Bessel beam raster scanning designs rely on spatial light-modulators (SLM) or axicons, and annular masks, which require additional optics to conjugate these components properly.

We do note that a two-photon Bessel beam likely can have a higher aspect ratio than a beam produced with a layered phase mask as presented here. The main difference between the two methods is that even a two-photon Bessel beam will produce pronounced sidelobes, whereas the phase mask presented here has a smooth, non-oscillatory transition into the background.

Bessel beams have also been hailed as being more robust to aberrations and optical occlusions. We do not know if some of these properties are also inherited for our EDF beam design. Since the effective focus is the superposition of many foci, which in turn each are essentially Bessel beams, it can be stipulated that such an arrangement is more robust to aberrations than interfering the whole wavefront coherently. We see another advantage for our method over Bessel beams in terms of adaptive optics. As our method uses the full pupil of the objective (in contrast, a Bessel beam only occupies a thin annulus), our method is expected to be compatible with adaptive optics that commonly use deformable mirrors conjugated to the pupil plane.

Lastly, it has been demonstrated that splitting an ultrafast laser beam into multiple, time-delayed beams can reduce the effects of photo-bleaching [11]. In that regard, our method represents a simple way to multiplex the laser pulse into five individual beams that arrive at slightly different times. As such, it would be expected that this could reduce photo-bleaching by a similar effect as previously published.

Overall, we present an unexpected use of a phase mask for ultrafast optics that was originally intended for linear optics using low coherence fluorescence light. We find that for the nonlinear application, the method has some advantages over other EDF approaches, such as high light-throughput, lack of sidelobes and being achromatic. We also think that due to its simplicity, this method will rapidly find widespread applications in multiphoton laser scanning, as existing instruments can be readily retrofitted for EDF imaging.

## 5.2 Acknowledgments

The authors would like to thank Vladimir Zhemkov and Hua Zhang for providing the fixed brain slice.

## Supplementary Information

### Supplementary notes

#### Supplementary Note 1

We further investigated how the EDF mask would perform using a higher NA of the objective was used for illumination. To this end, the telescope relaying the EDF mask to the galvo was changed to lenses with focal lengths of 125 mm and 100 mm, which results in an excitation NA of ~0.9.

Supplementary Figure 1 shows the resulting PSF with and without the EDF mask and the corresponding low NA Gaussian beam that matches the axial length of the EDF PSF. The Full width Half Maximum (FWHM) in the axial direction was 2.06 ± 0.13 μm for the full NA PSF and 10.17 ± 0.39 μm for the EDF beam. The lateral FWHM increased form 0.42 ± 0.03 μm (full NA) to 0.63 ± 0.04 μm for the EDF PSF. The low NA Gaussian has a lateral FWHM of 0.90 ± 0.04 μm.

#### Supplementary Note 2

A significant simplification for retrofitting an existing two photon raster scanning microscope would be achieved by simply placing the EDF mask right next to the galvanometric mirror. As such, the mask and this particular galvo cannot be perfectly conjugated to the pupil plane. However, we experimentally could not see much difference in the PSF when we slightly moved our EDF mask axially, so there is some robustness for axial misalignment of the mask. We thus removed the telescope that relays the EDF mask to the first galvo mirror and placed the mask as close as possible to the galvo.

Since the mask is larger than the diameter than the clear aperture of our galvanometric mirror, only ~four zones of the mask could be used. The FWHM in the axial direction was 1.49± 0.03 μm for the full NA PSF and 6.02 ± 0.09 μm for the EDF beam. The lateral FWHM increased form 0.38 ± 0.02 μm (full NA) to 0.55 ± 0.02μm for the EDF PSF. Still we observe the expected ~four-fold depth of focus increase in the PSF (Supplementary Figure 2). As such, we expect that a suitably sized EDF mask, which fits the clear aperture of the galvo mirror, can be directly used without an additional relay optics system.

We note that in this configuration, the entire NA of the objective was used, which is a regime where we found the EDF mask not to operate most effectively.

To use an effective smaller excitation NA as presented in the main manuscript, two approaches seem feasible. Either a smaller EDF mask would need to be used to match the existing microscope, or larger galvo mirrors, together with a smaller magnification to the pupil plane, would need to be used.

### Supplementary Figures

**Supplementary Fig. 1.**
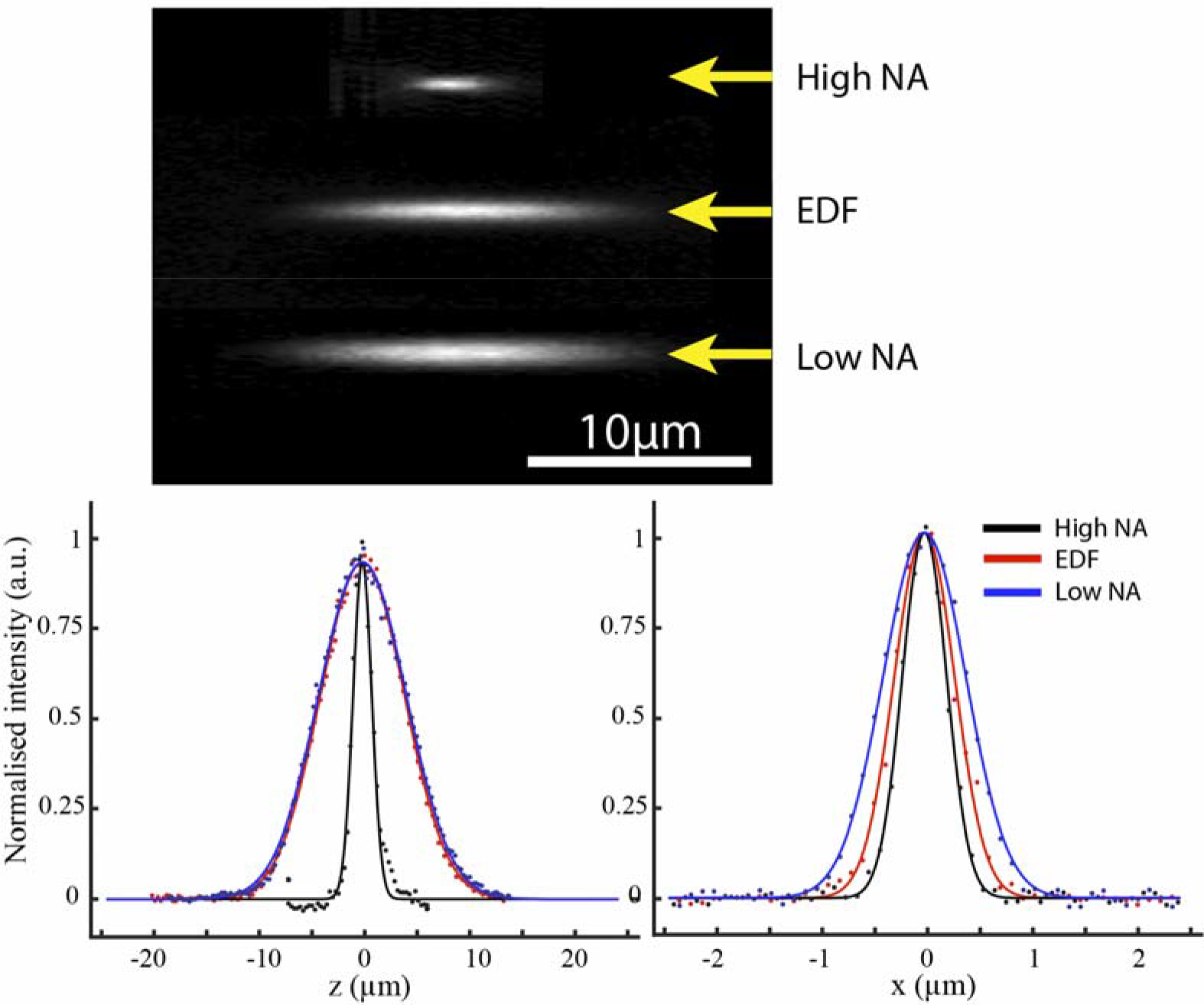
Point spread functions measured with 200nm green fluorescent beads using an NA of 0.9. Top panel, from top to bottom: PSF for laser focus using full NA of 0.0 and no EDF mask. PSF using EDF phase mask placed directly in front of the galvo mirror (no relay telescope).: Axial and lateral profiles through the point spread functions. Black curve corresponds to full NA PSF without EDF mask, red curve corresponds to PSF using the EDF mask and blue curve corresponds to low NA laser focus.

**Supplementary Fig. 2.**
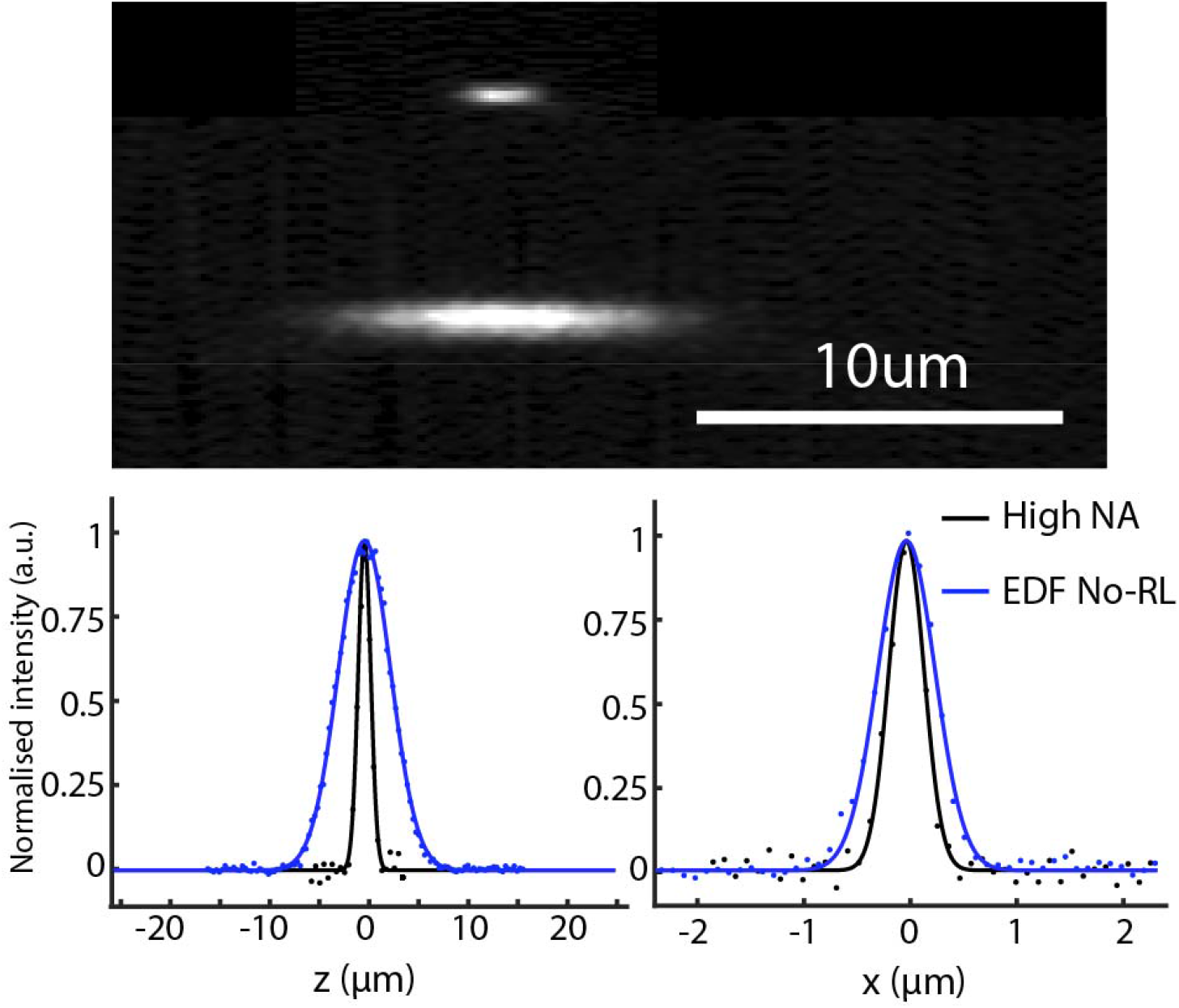
Point spread functions measured with 200nm green fluorescent beads. Top panel, from top to bottom: PSF for laser focus using full NA and no EDF mask. PSF using EDF phase mask placed directly in front of the galvo mirror (no relay telescope).: Axial and lateral profiles through the point spread functions. Black curve corresponds to full NA PSF without EDF mask, red curve corresponds to PSF using the EDF mask and blue curve corresponds to low NA laser focus.

